# The Axin scaffold protects the kinase GSK3β from cross-pathway inhibition

**DOI:** 10.1101/2022.12.05.519208

**Authors:** Maire Gavagan, Noel Jameson, Jesse G. Zalatan

## Abstract

Multiple signaling pathways regulate the kinase GSK3β by inhibitory phosphorylation at Ser9, which then occupies the GSK3β priming pocket and blocks substrate binding. Since this mechanism should affect GSK3β activity towards all primed substrates, it is unclear why Ser9 phosphorylation does not affect other GSK3β-dependent pathways, such as Wnt signaling. We used biochemical reconstitution and cell culture assays to evaluate how Wnt-associated GSK3β is insulated from cross-activation by other signals. We found that the Wnt-specific scaffold protein Axin allosterically protects GSK3β from phosphorylation at Ser9 by upstream kinases, which prevents accumulation of pS9-GSK3β in the Axin-GSK3β complex. Scaffold proteins that protect bound proteins from alternative pathway reactions could provide a general mechanism to insulate signaling pathways from improper crosstalk.

## Introduction

Glycogen Synthase Kinase 3β (GSK3β) is a potential therapeutic target for a range of diseases (Beurel et al., 2015; Nusse and Clevers, 2017), but targeting GSK3β is complicated because it has important roles in multiple signaling pathways (Bhat et al., 2018). Understanding how GSK3β is regulated by different signaling pathways could enable strategies to target distinct sub-populations of GSK3β.

Both Wnt and growth factor/insulin signaling pathways regulate GSK3β, but these pathways do not cross-activate (Ding et al., 2000; McManus et al., 2005; Ng et al., 2009). In Wnt signaling, the scaffold protein Axin binds GSK3β, its substrate β-catenin, and other proteins in a Wnt-specific complex called the destruction complex. Wnt signals inhibit kinase reactions in this complex (Hernández et al., 2012; Stamos et al., 2014), providing a mechanism to regulate Wnt-associated GSK3β without affecting other GSK3β-dependent pathways (Beurel et al., 2015; Gavagan et al., 2020). In contrast, in growth factor/insulin signaling, the kinases PKA and PKB/Akt phosphorylate GSK3β at Ser9 (Cross et al., 1995; Fang et al., 2000; Jensen et al., 2007; Sutherland et al., 1993), which inhibits GSK3β by binding in the priming pocket and blocking substrate binding (Dajani et al., 2001; Frame and Cohen, 2001; Stamos et al., 2014; ter Haar et al., 2001) (Figure 1A). It remains unclear why growth factor/insulin signaling does not globally inhibit GSK3β and cross-activate the Wnt pathway.

**Figure 1.**
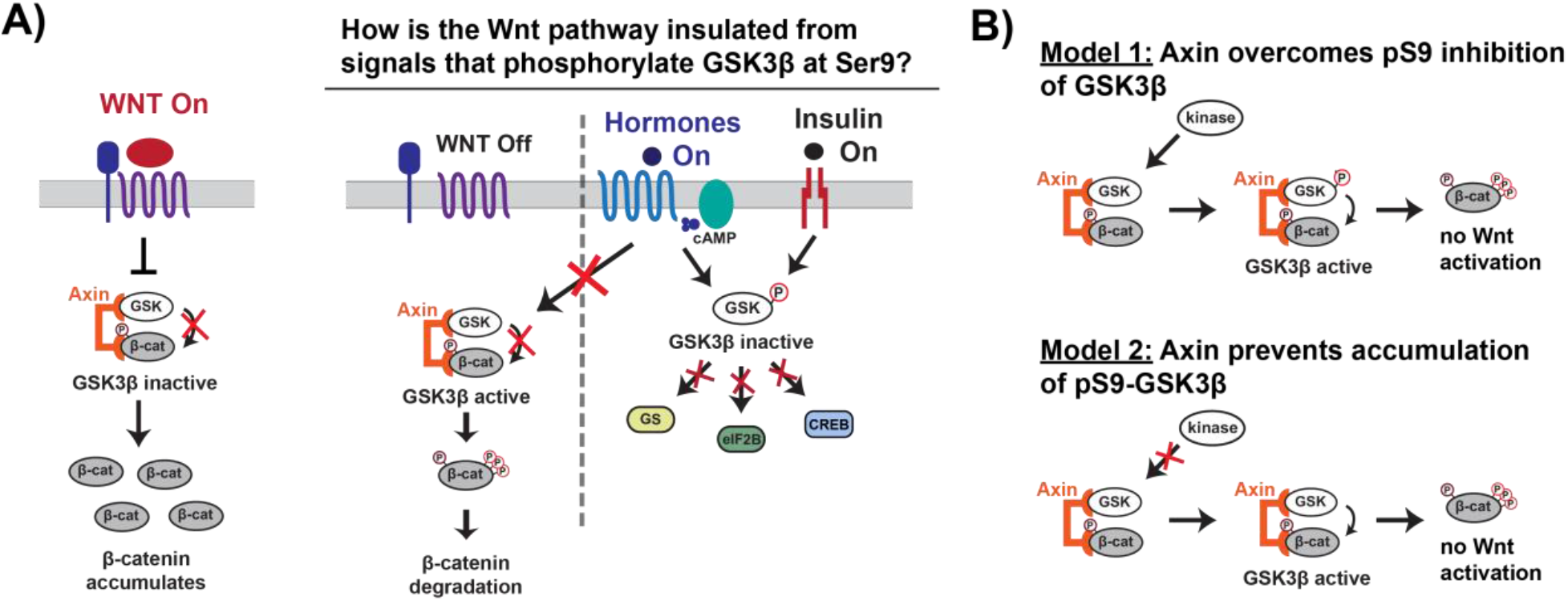
Wnt signaling is insulated from signals that phosphorylate GSK3β at Ser9. A) In the Wnt pathway, the scaffold protein Axin coordinates a GSK3β complex that phosphorylates β-catenin, which is then degraded. Wnt signals inhibit phosphorylation, allowing β-catenin levels to rise and initiate a transcriptional program (Nusse and Clevers, 2017). In other signaling pathways, upstream signals regulate GSK3β through phosphorylation at Ser9, which blocks substrate binding, inhibits activity towards downstream substrates, and activates downstream signaling (Sutherland, 2011). B) The scaffold protein Axin could insulate Wnt-associated GSK3β from Ser9 inhibition by restoring GSK3β activity towards β-catenin even when phosphorylated at Ser9 (Model 1) or by preventing accumulation of pS9-GSK3β in the Wnt destruction complex (Model 2).

Previous work in the field suggests two potential biochemical mechanisms that could insulate Wnt signaling from insulin and growth factor signals. First, by tethering GSK3β and the Wnt substrate β-catenin together, the Axin scaffold could rescue enzyme activity from the inhibitory effects of Ser9 phosphorylation (Figure 1B) (Beurel et al., 2015; Frame and Cohen, 2001). A second possibility is that Axin prevents accumulation of pS9-GSK3β, either through direct steric effects or indirect allosteric effects (Figure 1B). This model is supported by *in vivo* experiments showing that in insulin-treated cells, Ser9 phosphorylation increases in the total GSK3β population but is unchanged in the Axin-associated GSK3β pool (Ding et al., 2000; Ng et al., 2009). Using a reconstituted biochemical system, we found that Axin allosterically protects GSK3β from phosphorylation at Ser9. The ability of scaffold proteins to allosterically regulate bound enzymes and substrates is well-established (Good et al., 2011), but the use of similar mechanisms to prevent competing, scaffold-independent signaling reactions has not previously been characterized. Our findings suggest a new mechanism for how scaffold proteins can shield bound proteins to promote specificity in interconnected signaling networks.

## Results and Discussion

### Phosphorylation at Ser9 inhibits GSK3β

It is well-established that Ser9 phosphorylation inhibits GSK3β activity, but quantitative measurements are limited and variable (Frame and Cohen, 2001; Stambolic and Woodgett, 1994; Sutherland et al., 1993). To assess if the Wnt pathway can overcome Ser9 phosphorylation, we need quantitative metrics for comparison. We therefore used a biochemically-reconstituted system to measure initial rates for the GSK3β reaction with pS45-β-catenin and determined the steady state kinetic parameters *k*_cat_, *K*_M_, and *k*_cat_/*K*_M_ as described previously (Gavagan et al., 2020). We used PKA to prepare fully-phosphorylated pS9-GSK3β (see Methods and Figure S1C). We observed that phosphorylation of GSK3β at Ser9 decreases *k*_cat_/*K*_M_ towards pS45-β-catenin by a factor of ∼200-fold compared with unphosphorylated or mutant S9A GSK3β (Figure 2B). The observed rates for unphosphorylated GSK3β and PKA-treated GSK3β_S9A are indistinguishable, indicating that the large rate decrease in pS9-GSK3β is from phosphorylation at Ser9, not any other unknown PKA phosphosites. pS9-GSK3β does not detectably saturate at high substrate concentration, giving a limit for the *K*_M_ of ≥2 μM. The >7-fold increase in *K*_M_ is consistent with the model that Ser9 phosphorylation inhibits GSK3β by interfering with substrate binding, although we cannot rule out the possibility that pSer9 also affects other catalytic steps.

**Figure 2.**
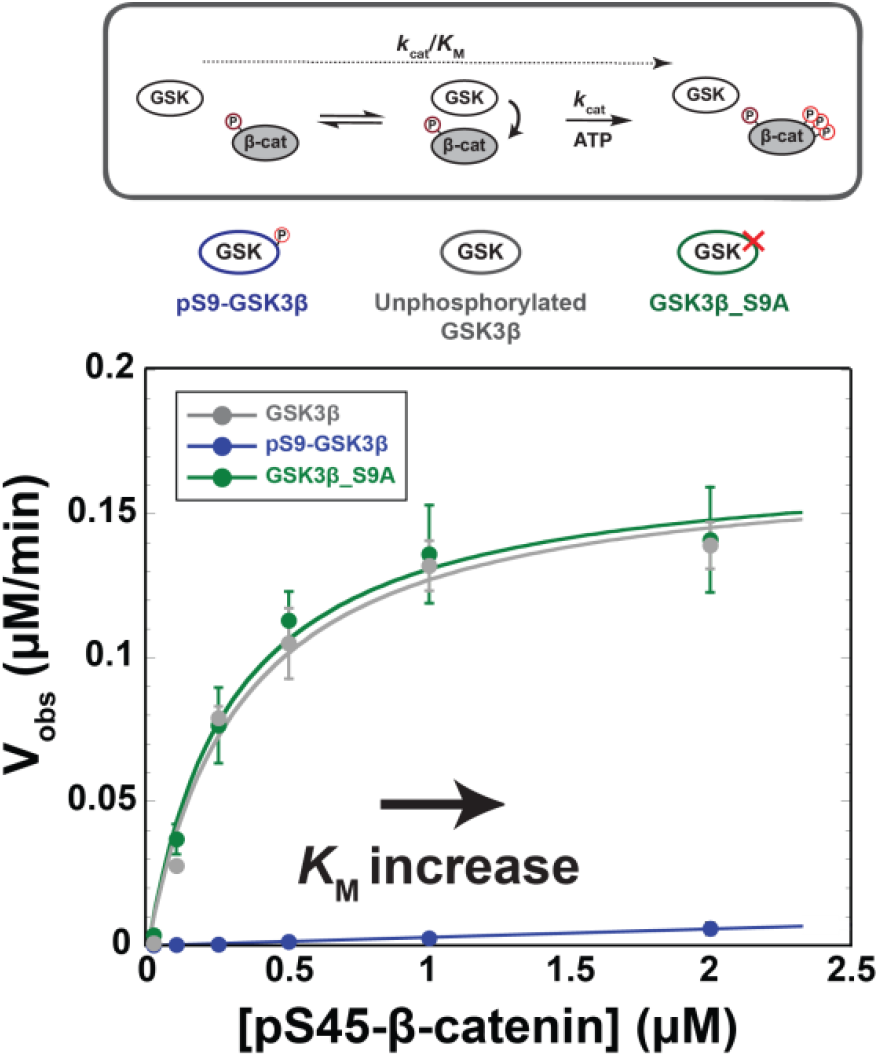
Phosphorylation at Ser9 inhibits GSK3β activity towards pS45-β-catenin. Kinetic scheme and Michaelis-Menten plots for reactions of unphosphorylated GSK3β, pS9-GSK3β, and GSK3β_S9A with pS45-β-catenin. Plots are *V*_obs_ versus [pS45-β-catenin] at 10 nM GSK3β. GSK3β phosphorylates pS45-β-catenin at three sites: S33, S37, and T41. Values are mean ± SD for at least 3 measurements. See Table S1 for values of fitted kinetic parameters.

### The scaffold protein Axin cannot overcome pS9-GSK3β inhibition

Addition of Axin to reactions with unphosphorylated GSK3β and PKA-treated GSK3β_S9A produced modest ∼2-fold increases in *k*_cat_/*K*_M_ arising from small changes to both *k*_cat_ and *K*_M_, (Figure 3), consistent with previous results (Gavagan et al., 2020). In the pS9-GSK3β reaction, however, Axin produced a ∼20-fold *k*_cat_/*K*_M_ increase. Notably, this effect is primarily due to a decrease in the *K*_M_ to 0.27 µM, similar to the values for unphosphorylated GSK3β and GSK3β_S9A (Figure 3A). This result suggests that Axin can compensate for the inhibitory effect of pS9-GSK3β on substrate binding, possibly because the Axin binding site for β-catenin allows formation of an Axin•GSK3β•β-catenin ternary complex even when the GSK3β priming pocket is blocked.

**Figure 3.**
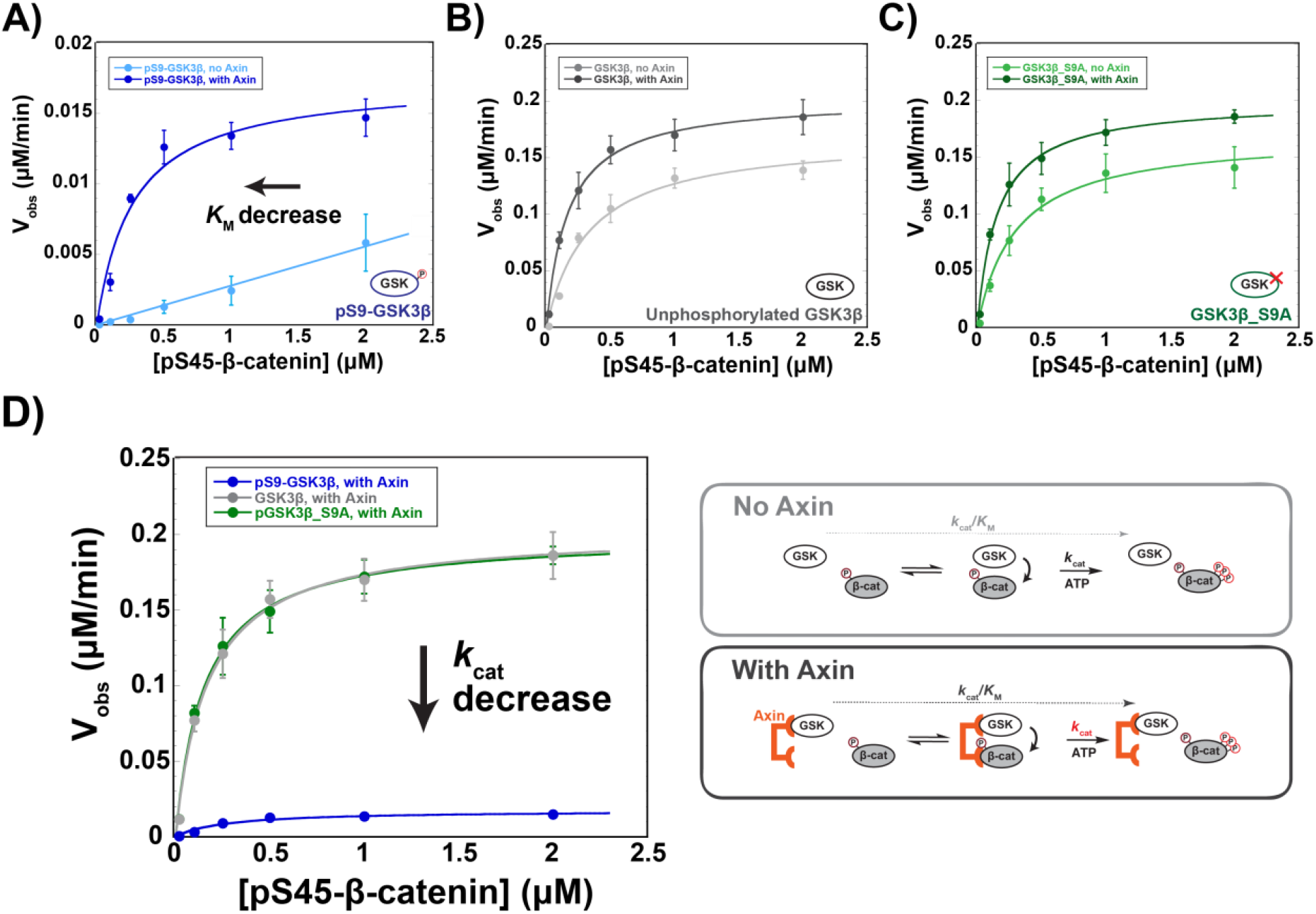
Axin restores the *K*_M_ for β-catenin but cannot overcome pS9-GSK3β Inactivation. A-C) Michaelis-Menten plots of *V*_obs_ versus [pS45-β-catenin] in the presence and absence of 500 nM Axin with 10 nM pS9-GSK3β (A), unphosphorylated GSK3β (B) or PKA-treated GSK3β_S9A (C). At the Axin concentrations used in these experiments all the GSK3β is bound to Axin (Gavagan et al., 2020). D) Minimal kinetic scheme and Michaelis-Menten plots for reactions of GSK3β with pS45-β-catenin in the presence of Axin plotted on the same scale. Values are mean ± SD for at least 3 measurements. See Table S1 for values of fitted kinetic parameters.

Although Axin appears to fully rescue the *K*_M_ effect from Ser9 phosphorylation, there is still a substantial ∼10-fold *k*_cat_ decrease. The simplest interpretation of this result is that Axin assembles a non-productive pS9-GSK3β•pS45-β-catenin complex that is still inhibited by pSer9 occupying the priming pocket. Thus, if pS9-GSK3β accumulates in the Axin-mediated destruction complex, β-catenin phosphorylation will be inhibited by ∼10-fold, likely leading to improper activation of the Wnt pathway. Stimulation with high levels of Wnt ligand produces ∼5-fold decreases in GSK3β phosphorylation of β-catenin (Hannoush, 2008; Hernández et al., 2012), and *in vivo* changes in β-catenin levels as low as 2-fold can have measurable effects on transcription of Wnt output genes (Jacobsen et al., 2016).

### Axin prevents accumulation of pS9-GSK3β in the destruction complex

To test if Axin-bound GSK3β is shielded from upstream kinases, we evaluated the effect of Axin on PKA, a kinase upstream of GSK3β in growth factor signaling (Fang et al., 2000). We found that Axin produced a 7-fold drop in *k*_cat_/*K*_M_ for PKA phosphorylation of GSK3β at Ser9, primarily from a 4-fold increase in *K*_M_ (Figure 4A). This *K*_M_ increase suggests that Axin interferes with formation of the PKA•GSK3β complex. To determine if this effect is specific to GSK3β, we measured the effect of Axin on the reaction with another PKA substrate, CREB (Naqvi et al., 2014). Axin has no effect on observed rates or *k*_cat_/*K*_M_ for the CREB reaction (Figure 4B). This result indicates that Axin is not a competitive inhibitor of PKA at the concentrations used in our assays, nor is Axin directly binding PKA to regulate its activity.

**Figure 4.**
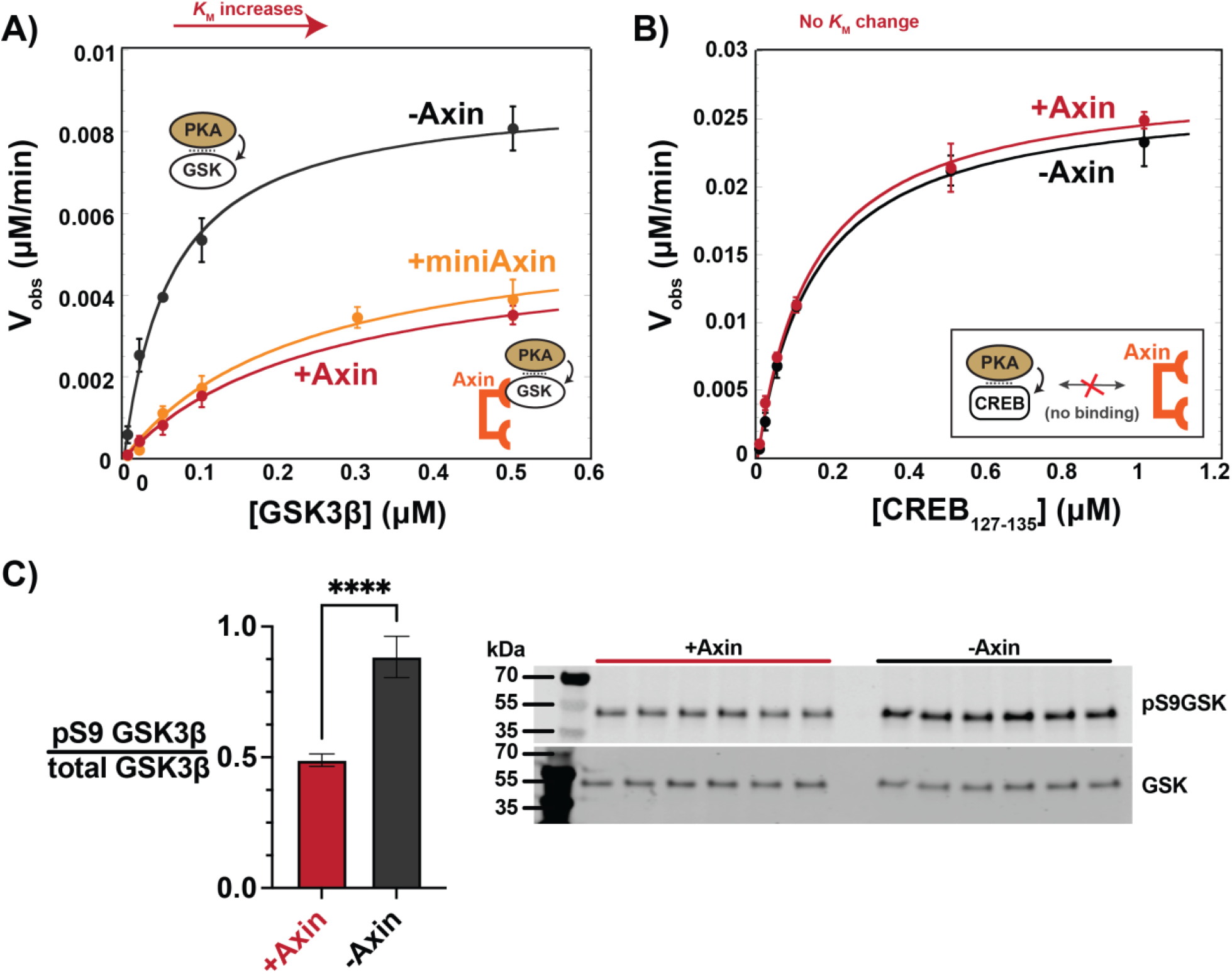
Axin prevents phosphorylation of GSK3β at Ser9. A) Michaelis-Menten plots of *V*_obs_ versus [GSK3β] with 20 nM PKA in the presence and absence of 500 nM Axin. B) Michaelis-Menten plots of *V*_obs_ versus [CREB_127-135_] with 20 nM PKA in the presence and absence of 500 nM Axin. Values are mean ± SD for at least 3 measurements. See Table S1 for values of fitted kinetic parameters from (A) and (B). C) Normalized Western blot analysis and blot images of pS9-GSK3β in HEK293 cells transiently expressing Axin or a negative control (see Methods). Normalized pS9-GSK3β levels were calculated by dividing pS9-GSK3β signal by total GSK3β and averaging across six independent experiments. The p-value between Axin-expressing cells and a non-Axin negative control is <0.0001.

There are several plausible models for how the Axin•GSK3β interaction might disrupt PKA phosphorylation. The simplest model is that Axin sterically occludes upstream kinases from accessing the Ser9 site on GSK3β. The Axin binding site on GSK3β does not directly overlap with Ser9, nor is it immediately adjacent (Ikeda et al., 1998), but Axin is a large, disordered protein and could potentially extend towards the N-terminus of GSK3β. Another possibility is that Axin binding to GSK3β produces allosteric changes that make it less accessible to upstream kinases.

To distinguish between these possible models, we measured the effect of a minimal Axin truncation on PKA phosphorylation of GSK3β. This miniAxin fragment (residues 384-518) binds GSK3β with a similar affinity as full-length Axin_1-826_ (Gavagan et al., 2020). Because it is substantially smaller than full-length Axin, miniAxin is less likely to sterically block PKA from accessing GSK3β. Addition of miniAxin produced a 6-fold decrease in *k*_cat_/*K*_M_ with a 4-fold increase in *K*_M_, similar to full length Axin (Figure 4A). This result supports the model that Axin binding leads to allosteric changes in GSK3β that make it less accessible to PKA. However, we cannot definitively rule out the possibility that miniAxin has a steric effect on PKA phosphorylation of GSK3β.

To determine if Axin prevents pS9-GSK3β accumulation *in vivo*, we overexpressed Axin in HEK293 cells. Consistent with previous results, pS9-GSK3β can be detected in cells grown in serum-containing media (Fang et al., 2000; Whiting et al., 2015). When Axin is overexpressed, we observed a significant decrease in pS9-GSK3β levels (Figure 4C), consistent with our observation that Axin prevents phosphorylation of Ser9 *in vitro*.

## Conclusions

The observation that Axin protects GSK3β from PKA phosphorylation *in vitro* is consistent with previous *in vivo* co-immunoprecipitation experiments suggesting that Axin-associated GSK3β is not phosphorylated at Ser9 (Ng et al., 2009). Beyond Axin-mediated shielding of GSK3β, other mechanisms could also contribute to preventing accumulation of pS9-GSK3β in the Wnt destruction complex. Axin interacts with the phosphatase PP2A and may promote PP2A-mediated dephosphorylation of pS9-GSK3β (Cantoria et al., 2022). Alternatively, sub-cellular localization or phase separation could sequester GSK3β in distinct pools that are associated with different signaling pathways and independently regulated (Anton et al., 2022; Bock et al., 2020; Su et al., 2016; Zhang et al., 2020). Wnt pathway proteins also phase separate (Nong et al., 2021; Schaefer et al., 2018), which could exclude kinases like Akt and PKA from accessing Wnt-associated GSK3β. Although these other mechanisms may play an important role, here we have used biochemical reconstitution to systematically evaluate two possible direct contributions of the Axin scaffold to pathway insulation, and our results suggest that Axin can allosterically control the accessibility of GSK3β to upstream signals from competing pathways.

## Acknowledgements

We thank Dustin Maly, John Scott, Maryanne Kihiu and members of the Zalatan group for comments and discussion. This work was supported by NIH R35 GM124773 (J.G.Z.).

## Author contributions

M.G, N.J., and J.G.Z. designed research; M.G and N.J. performed research; and M.G, N.J., and

J.G.Z. wrote the paper.

## Abbreviations

GSK3β: Glycogen synthase kinase 3β
CREB: cAMP response element binding protein
PKA: protein kinase A.

