## Supplementary Information for "The Axin scaffold protects the kinase GSK3β from cross-pathway inhibition"

This file contains:

Materials and Methods
Supplemental Figures 1-12
Supplemental Tables 1-6
Supplemental References

### Materials and Methods

#### *Protein Expression Constructs*

The proteins Axin, pS45-β-catenin, PKA, and the $CREB_{127-135}$ peptide ILSRRPSYR were cloned and expressed as previously described (Gavagan et al., 2020). All sequences except PKA were cloned into *E. coli* expression vectors containing an N-terminal maltose binding protein (MBP) and a C-terminal His6 tag. The catalytic subunit of mouse PKA was expressed from pET15b with an N-terminal His-tag (addgene #14921)(Narayana et al., 1997).

pS45-β-catenin was produced by coexpression with CK1α, as previously described (Gavagan et al., 2020). Lambda phosphatase (λPPase) was cloned with an N-terminal GST tag and a C-terminal His6 tag; the human λPPase sequence was obtained from VMG950 (Good et al., 2009). Dephosphorylated GSK3β was produced by coexpression with λPPase. The coexpression plasmid for GSK3β and λPPase was constructed by inserting the GST-λPPase expression cassette (without the His6 tag) into the MBP-GSK3β-His6 plasmid. GSK3β point mutants and Axin truncations were constructed by assembling PCR fragments. Unless otherwise noted, all wt and mutant GSK3β constructs in this work were coexpressed with λPPase to ensure they are unphosphorylated.

#### *Protein Expression and Purification*

For quantitative kinetic and binding assays, all Wnt pathway, CREB, and PKA proteins were expressed in Rosetta (DE3) pLysS *E. coli* cells by inducing with 0.5 mM IPTG overnight at 18 ºC. Constructs with N-terminal MBP and C-terminal His6 tags (pS45-β-catenin, Axin, and $CREB_{127-135}$) were affinity purified with HisPur Ni-NTA resin (Thermo Scientific) and amylose resin (NEB). The PKA catalytic subunit was purified on Ni-NTA resin. pS45-β-catenin was produced by coexpression with CK1α, as previously described (Gavagan et al., 2020).

Unphosphorylated GSK3β and GSK3β_S9A were produced by coexpression with lambda phosphatase and affinity purified with HisPur Ni-NTA resin (Thermo Scientific). Treatment of GSK3β and GSK3β_S9A with lambda phosphatase produces an ~5-fold increase in $k_{cat}/K_M$ that is due to an ~5-fold increase in $k_{cat}$ (Figure S12 & Tables S4 & S5). To produce PKA-treated pS9-GSK3β and GSK3β_S9A, 10 μM of λPPase-treated, Ni-NTA-purified GSK3β and GSK3β_S9A were incubated with 5 μM PKA and 500 μM ATP for 2 hrs at 25 °C. Phosphorylated GSK3β and GSK3β_S9A were separated from PKA by affinity purification with amylose resin (NEB). GSK3β phosphorylation at Ser9 and Tyr216 was assessed by western blot using antibodies for pSer9 GSK3β (Cell Signaling Technology #9561), pTyr216 GSK3β (BD Biosciences #612312), and MBP (Cell Signaling Technology #2396) (Figure S1). The secondary antibodies were IRDye 800CW Goat Anti-Rabbit IgG antibody (Li-Cor #926-32211) for pSer9 and pTyr216 GSK3β and IRDye 800CW Donkey Anti-Mouse IgG antibody (Li-Cor #926-32212) for MBP.

Purified proteins were dialyzed into 20 mM Tris-HCl pH 8.0, 150 mM NaCl, 10% glycerol, and 2 mM DTT at 4 °C, aliquoted and stored at -80 °C. If necessary, proteins were concentrated using 10000 or 30000 MWCO Amicon Ultra-15 Centrifugal Filter devices at 4 °C, 2000×g. Protein concentrations were determined using a Bradford assay (Thermo Scientific). pS45-β-catenin was further purified by size exclusion chromatography using a Superdex 200 Increase 10/300 GL column (GE Healthcare) to remove a copurifying fragment before being dialyzed into 20 mM Tris-HCl pH 8.0, 150 mM NaCl, 10% glycerol, and 2 mM DTT, aliquoted, and stored at -80 °C. A Coomassie gel showing the purity of the proteins in this work is shown in Figure S1A.

*Quantitative Kinetic Assays*

*In vitro* kinetic assays were conducted in kinase assay buffer [40 mM HEPES pH 7.4, 50 mM NaCl, 10 mM $MgCl_2$, and 0.05% IGEPAL] at 25 ºC in 60 µL total volume. Reactions were initiated by adding ATP to a final concentration of 100 µM. This ATP concentration is saturating for all reactions (Figure S3 & Table S3). Reaction timepoints for initial rate kinetics were obtained at 10, 30, 60, and 90 seconds (pS45-β-catenin reactions with unphosphorylated GSK3β and GSK3β_S9A, Figures S4 & S5); 1, 2, 5, and 10 minutes (pS45-β-catenin reactions with pS9-GSK3β, Figures S6 & S7); and 0.5, 1, 2, and 4 minutes (vary [GSK3β] and [$CREB_{127-135}$] reactions with PKA, Figures S8 & S9). 10 µL aliquots were quenched by boiling in 5X SDS loading buffer. Samples were analyzed by SDS-PAGE and quantitative western blotting as described below (Figures S4-S9). For reactions with pS45-β-catenin, all gel samples were diluted 5-fold in 1X SDS loading buffer to prevent a gel smearing artifact that occurs with [pS45-β-catenin] ≥ 500 nM. For reactions with PKA phosphorylation of GSK3β, samples were diluted 4-fold (500 nM GSK3β reactions without Axin) or 2-fold (all other GSK3β concentrations) to prevent signal saturation of the western blot scan.

GSK3β-phosphorylated β-catenin was detected using a primary anti-Phospho-β-Catenin (Ser33/37/Thr41) antibody (Cell Signaling Technology #9561) that recognizes triply phosphorylated pS33/pS37/pT41-β-catenin (Figure S2). PKA-phosphorylated GSK3β was detected using a primary anti-phospho-GSK3β (Ser9) antibody (Cell Signaling Technology #5558) that recognizes pS9-GSK3β (Figure S2). PKA-phosphorylated $CREB_{127-135}$ was detected using a primary anti-phospho-CREB (Ser133) antibody (Cell Signaling Technology #9198) that recognizes pS133-$CREB_{127-135}$ (Figure S2). For all reactions, the secondary antibody was IRDye 800CW Goat Anti-Rabbit IgG antibody (Li-Cor #926-32211).

Concentrations of phosphorylated product in each reaction were determined by comparing western blot signal intensities to an endpoint standard containing 50 nM product phosphorylated to completion (Figure S2). For pS45-β-catenin reactions the endpoint is pS45-β-catenin phosphorylated to completion by GSK3β as previously described (Gavagan et al., 2020). The pS45-β-catenin standard was prepared in a reaction with 50 nM pS45-β-catenin, 100 nM GSK3β, and 100 µM ATP in kinase assay buffer at 25 ºC for 15 min. For PKA phosphorylation of GSK3β reactions the endpoint is pS9-GSK3β, phosphorylated to completion by PKA. The pS9-GSK3β standard was prepared in a reaction with 50 nM dephosphorylated GSK3β, 100 nM PKA, and 500 µM ATP in kinase assay buffer at 25 ºC for 24 hours. To prevent signal saturation of the western blot scan, the pS9-GSK3β standard was diluted 4-fold in 1x SDS loading dye, to a final concentration of 12.5 nM pS9-GSK3β. For $CREB_{127-135}$ reactions the endpoint is pS133-$CREB_{127-135}$, phosphorylated to completion by PKA. The pS133-$CREB_{127-135}$ standard was prepared in a reaction with 50 nM $CREB_{127-135}$, 100 nM PKA, and 200 µM ATP in kinase assay buffer at 25 ºC for 20 hours.

Initial rate measurements were performed in triplicate. Phosphorylated product levels from quantitative western blots were analyzed using Image Studio Lite 5.2.5 (Li-Cor) and kinetic parameters were determined by fitting to the Michaelis-Menten equation or to a linear equation using Kaleidagraph 4.1.3. Initial rates for each reaction were determined by fitting a linear model to a graph of [product] vs time. Kinetic parameters were determined by fitting plots of initial rates ($V_{\text{obs}}$) vs. [substrate] to the Michaelis-Menten equation $V_{obs} = k_{\text{cat}}[\text{E}]_0[\text{S}]/(K_{\text{M}} + [\text{S}])$. Standard errors for $k_{\text{cat}}$ and $K_{\text{M}}$ reported in Tables 1 and 2 are from non-linear least squares fits to this equation. Standard errors for $k_{\text{cat}}/K_{\text{M}}$ were obtained by fitting an alternative form of the equation $V_{obs} = (k_{\text{cat}}/K_{\text{M}})[\text{E}]_0[\text{S}]/\big(1 + ([\text{S}]/K_{\text{M}})\big)$. For the pS9-GSK3β reactions without Axin, which did not detectably saturate, the value of $k_{cat}/K_M$ was obtained from the slope of linear fit to the plot of $V_{obs}$ vs. [substrate].

The reaction conditions for *in vitro* kinetics experiments were tested to confirm the underlying assumptions in the kinetic model. As expected, reaction rates increase linearly with increasing enzyme concentration in all reactions (Figure S10). We also identified the optimal scaffold concentration for all reactions (Figure S11). Scaffold-dependent reactions typically have optimal scaffold concentrations, and can be slow at high concentrations of scaffold protein when kinase and substrate are bound to different scaffolds (Gavagan et al., 2020; Levchenko et al., 2000) (Figure S11).

*Phos-tag gel analysis of GSK3β phosphorylation*

Phos-tag gels were prepared and run as previously described (Gavagan et al., 2020). After electrophoresis, the gel was incubated 3x with transfer buffer + 10 mM EDTA for 10 min before the transfer to increase transfer efficiency. Protein levels were detected with anti-MBP antibody (Cell Signaling Technology #2396) (Figure S12A). The secondary antibody was IRDye 800CW Donkey Anti-Mouse IgG antibody (Li-Cor #926-32212).

*In vivo cell culture experiments*

The Axin open reading frame (Gavagan et al., 2020) was cloned into the human expression vector pcDNA3.1(+) (Thermo Fisher) with a C-terminal mCherry tag. For a protein overexpression negative control, eGFP was cloned into the human expression vector pMAX (Lonza Bioscience).

HEK293 cells (ATCC #CRL-1573) were plated at $5 \times 10^5$ cells/mL in 10% FBS DMEM in a 24-well plate. 24 hours after plating, individual wells were transfected with 50 μL Opti-MEM media containing 1 μg DNA and 1.5 μL TurboFectin (Origene TF81001). 48 hours following transfection, cells were washed twice with 500 μL PBS and lysed on ice in 40 μL lysis buffer [20 mM TRIS pH 7.5, 30 mM NaCl, 20 mM NaF, 1% NP-40, 0.5% DOC, 0.1% SDS, HALT protease and phosphatase inhibitor mixture (Thermo Scientific #78440)]. Lysate was then centrifuged at 13,000×g for 10 minutes at 4 °C. The resulting supernatant was boiled for 10 minutes in 5X SDS loading buffer and analyzed by SDS page and quantitative western blotting.

pS9-GSK3β was detected using the same antibody used in kinetics assays (Cell Signaling Technology #5558). Total GSK3β was detected using a primary anti-GSK3β antibody (Cell Signaling Technology #9832). The secondary antibodies were IRDye 800CW Goat Anti-Rabbit IgG antibody (Li-Cor #926-32211) for pS9-GSK3β and IRDye 680RD Donkey Anti-Mouse IgG antibody for total GSK3β. pS9-GSK3β and total GSK3β concentrations were analyzed using Image Studio Lite 5.2.5 (Li-Cor).

### Supplemental Figures

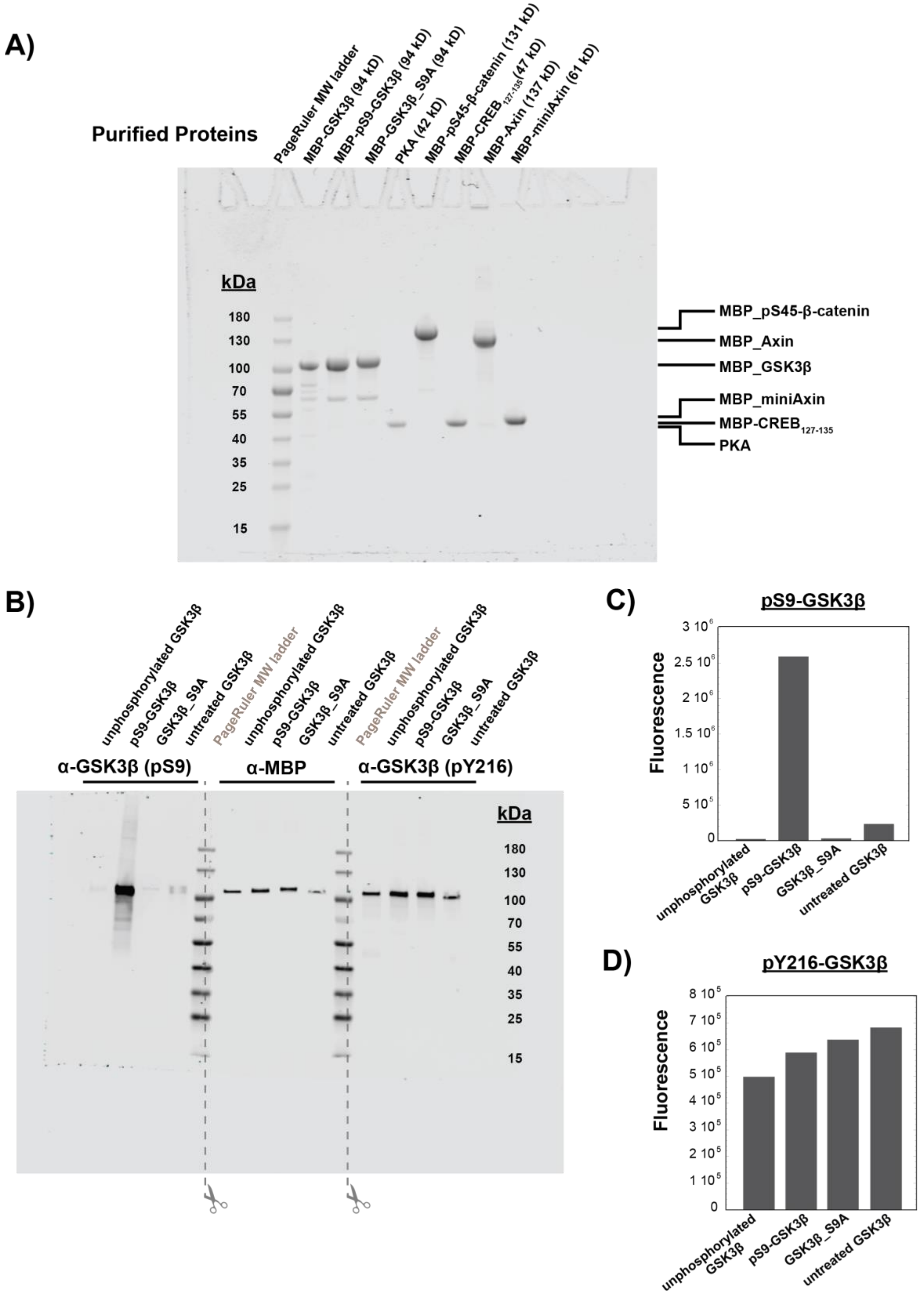

**Figure S1. Characterization of purified proteins, related to Figures 2-4**

A) Coomassie-stained SDS-PAGE of purified proteins used in this work. All proteins except PKA were purified as MBP fusion proteins (see Methods). Unphosphorylated GSK3β was purified after coexpression with lambda phosphatase (see Methods). pS45-β-catenin was purified after coexpression with CK1α as described previously (Gavagan et al., 2020). Phosphorylated GSK3β and GSK3β_S9A were purified after *in vitro* phosphorylation with PKA (see Methods). Each lane was loaded with 10 μL of 4 μM protein.

B) Western blot for phosphorylation state of GSK3β at Ser9 and Tyr216. GSK3β samples are unphosphorylated GSK3β, pS9-GSK3β, GSK3β_S9A, and untreated GSK3β (unmodified recombinant protein, not coexpressed with lambda phosphatase or treated with PKA). After the western blot transfer, the membrane was cut down the center of the MW ladder lanes so each third of the membrane could be incubated with separate antibodies (α-pS9-GSK3β, α-MBP for total protein, and α-pY216-GSK3β). The membrane fragments were placed back together for imaging.

C) PKA phosphorylates GSK3β at Ser9. The extent of Ser9 phosphorylation was quantified by western blot (Figure S1B). Fluorescence values were normalized using the α-MBP total protein loading control. No significant phosphorylation at Ser9 was detected for unphosphorylated GSK3β (phosphatase-treated) or GSK3β_S9A. Untreated GSK3β is partially (~10%) phosphorylated at Ser9.

D) Recombinant GSK3β is phosphorylated at Tyr216. The extent of Tyr216 phosphorylation was quantified by western blot (Figure S1B). Fluorescence values were normalized using the α-MBP total protein loading control.

A) Representative western blots for initial rate assays

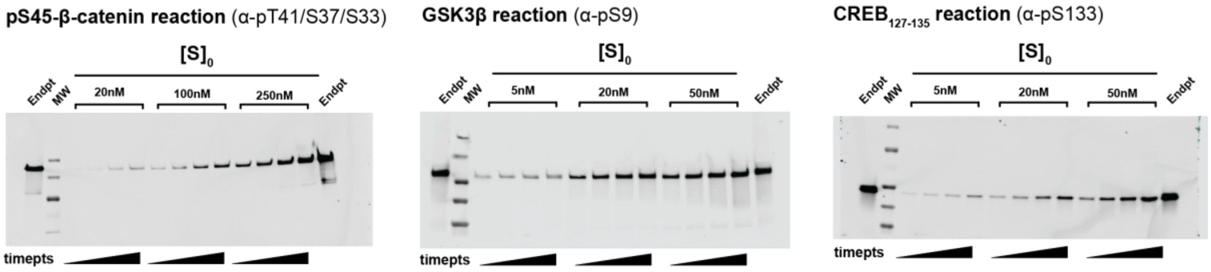


B) Validation of endpoint standards

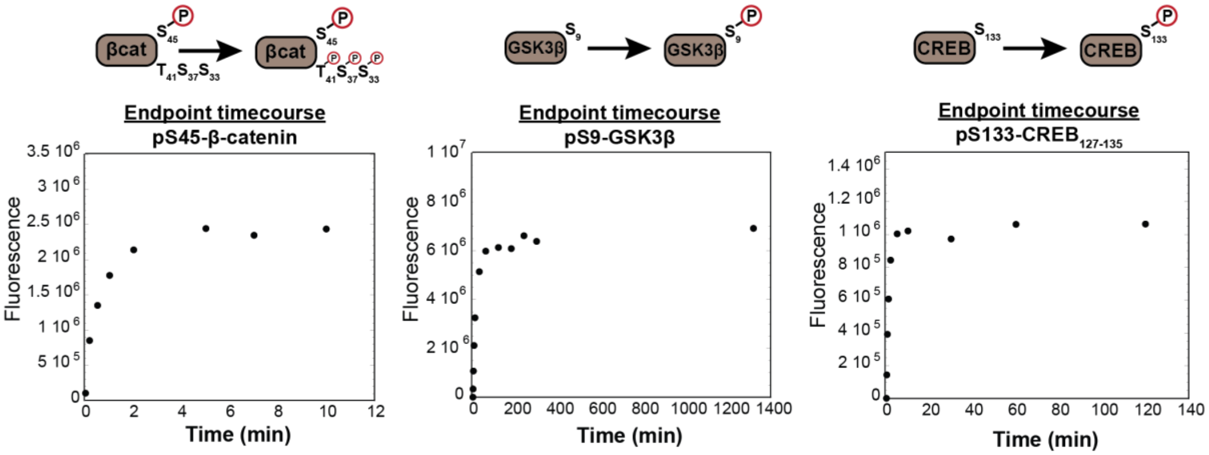


C) Antibody signal is linear

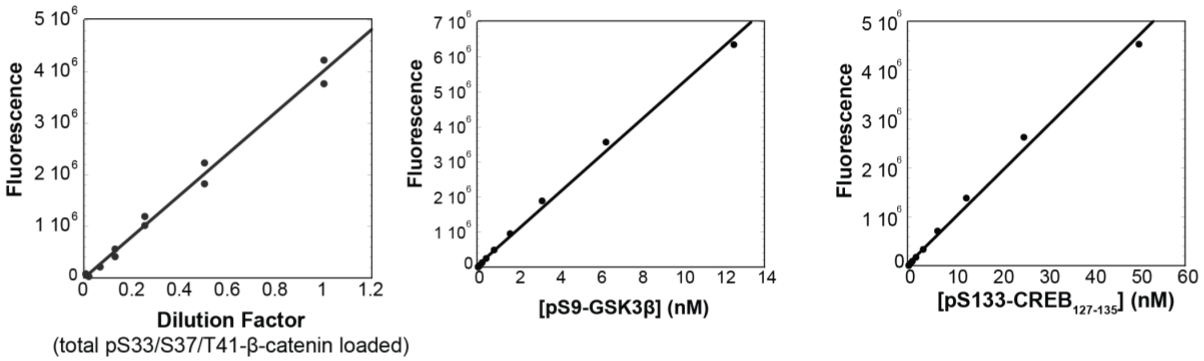

**Figure S2. Protein phosphorylation kinetic assays, related to Figures 2-4**

A) Representative western blots for reactions of GSK3β with pS45-β-catenin, PKA with GSK3β, and PKA with $CREB_{127-135}$. Reactions were conducted with 10 nM GSK3β or 20 nM PKA and the substrate concentrations indicated. Each gel was cut before transferring to the membrane to facilitate multiple simultaneous transfers in the same apparatus (as seen in Figures S4, S6, and S8). The images shown are the complete, uncropped blot membrane images.

B) Timecourses of phosphorylation of endpoint standards for GSK3β-phosphorylated pS45-β-catenin, PKA-phosphorylated pS9-GSK3β, and PKA-phosphorylated pS133-$CREB_{127-135}$.

C) The antibody signal is linear over a broad range spanning the observed signal in kinetic assays. The pS33/pS37/pT41-β-catenin data shows a set of 2-fold serial dilutions from a reaction with 3 µM pS45-β-catenin, 20 nM GSK3β and 500 nM miniAxin at the 1.5 minute timepoint. These data were published previously (Gavagan et al., 2020). The pS9-GSK3β and pS133- $CREB_{127-135}$ plots show sets of 2-fold serial dilutions from a 1:4 dilution of pS9-GSK3β endpoint (PKA reactions with GSK3β) or undiluted endpoint (PKA reactions with $CREB_{127-135}$), respectively.

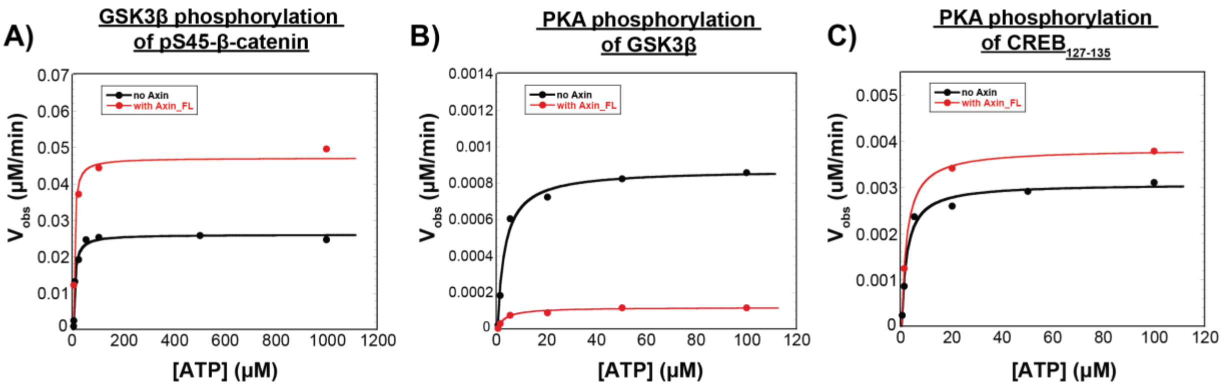


**Figure S3. The concentration of ATP used for quantitative kinetic experiments (100 μM) is saturating for all reactions, related to Figures 2-4.**

A) Michaelis-Menten plot of $V_{obs}$ vs. [ATP] at 10 nM unphosphorylated GSK3β and 50 nM pS45-β-catenin in the presence and absence of 500 nM Axin. Fits to the Michaelis-Menten equation give $K_{M,\ ATP}$ values of 4.6 ± 0.9 μM and 3.7 ± 1.3 μM in the presence and absence of Axin, respectively.

B) Michaelis-Menten plot of $V_{obs}$ vs. [ATP] at 20 nM PKA and 20 nM GSK3β in the presence and absence of 500 nM Axin. Fits to the Michaelis-Menten equation give $K_{M,\ ATP}$ values of 3.0 ± 0.6 μM and 3.3 ± 0.8 μM in the presence and absence of Axin, respectively.

C) Michaelis-Menten plot of $V_{obs}$ vs. [ATP] at 20 nM PKA and 20 nM $CREB_{127-135}$ in the presence and absence of 500 nM Axin. Fits to the Michaelis-Menten equation give $K_{M,\ ATP}$ values of 2.1 ± 0.4 μM and 2.1 ± 0.2 μM in the presence and absence of Axin, respectively.

$K_M$ values for (A)-(C) are compiled in Table S3.

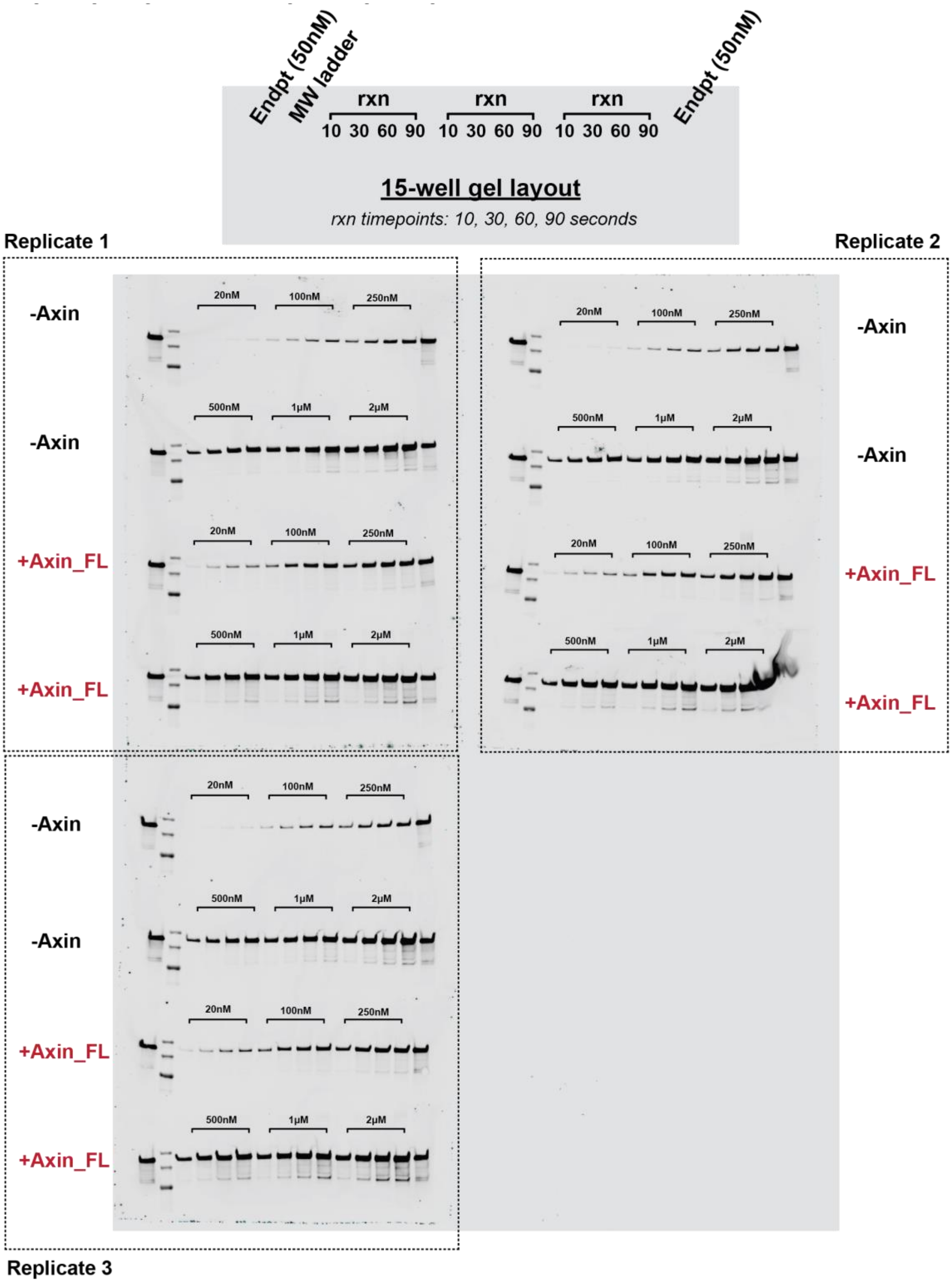


**Figure S4. Representative western blots for the reaction of unphosphorylated GSK3β with pS45-β-catenin in the presence and absence of Axin, related to Figure 2.**

Western blots for reactions of varying concentrations of pS45-β-catenin with 10 nM GSK3β in the presence and absence of 500 nM Axin. All gel samples were diluted 1:5 to prevent a gel smearing artifact (see Methods). See Figure S5 for quantification.

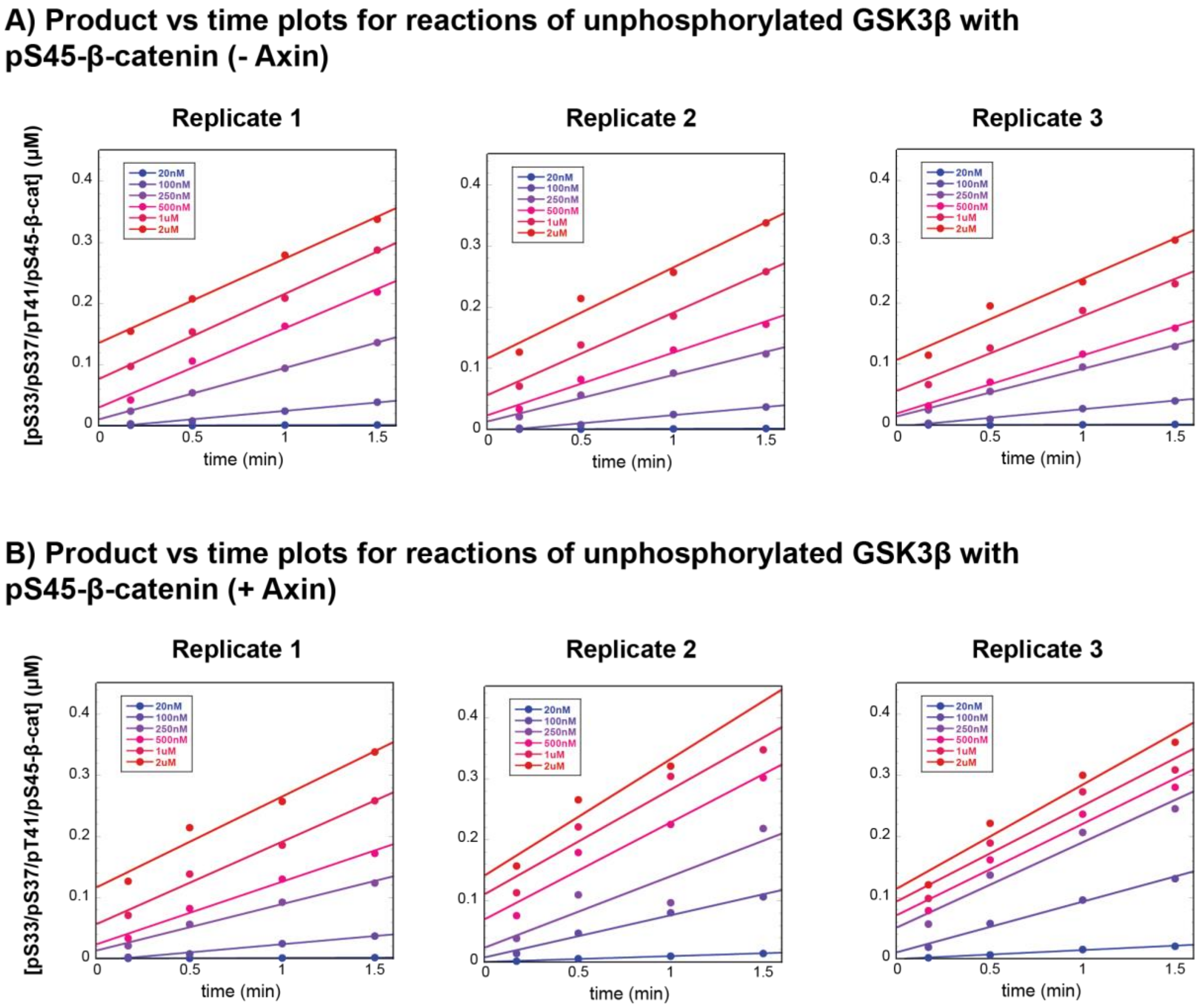


**Figure S5. Plots of product vs. time for reaction of GSK3β with pS45-β-catenin in the presence and absence of Axin, related to Figures 2 and 3.**

A) Product vs. time plots for reactions of unphosphorylated GSK3β with pS45-β-catenin in the absence of Axin.

B) Product vs. time plots for reactions of GSK3β with pS45-β-catenin in the presence of 500 nM Axin. Data in (A) and (B) correspond to the reaction conditions and western blots shown in Figure S4.

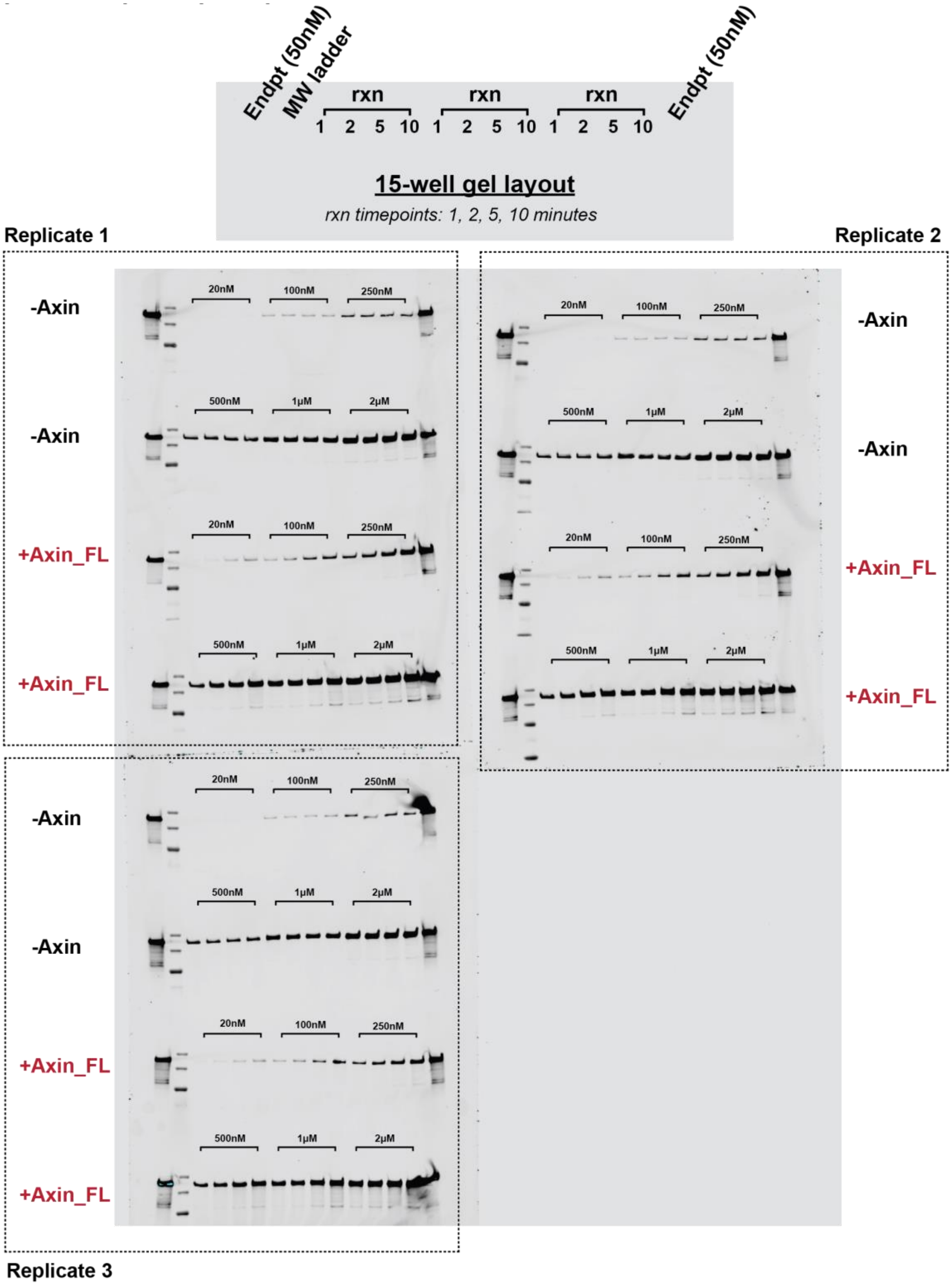


**Figure S6. Representative western blots for the reaction of pS9-GSK3β with pS45-β-catenin in the presence and absence of Axin, related to Figure 2.**
Western blots for reactions of varying concentrations of pS45-β-catenin with 10 nM pS9-GSK3β in the presence and absence of 500 nM Axin. All gel samples were diluted 1:5 to prevent a gel smearing artifact (see Methods). See Figure S7 for quantification.

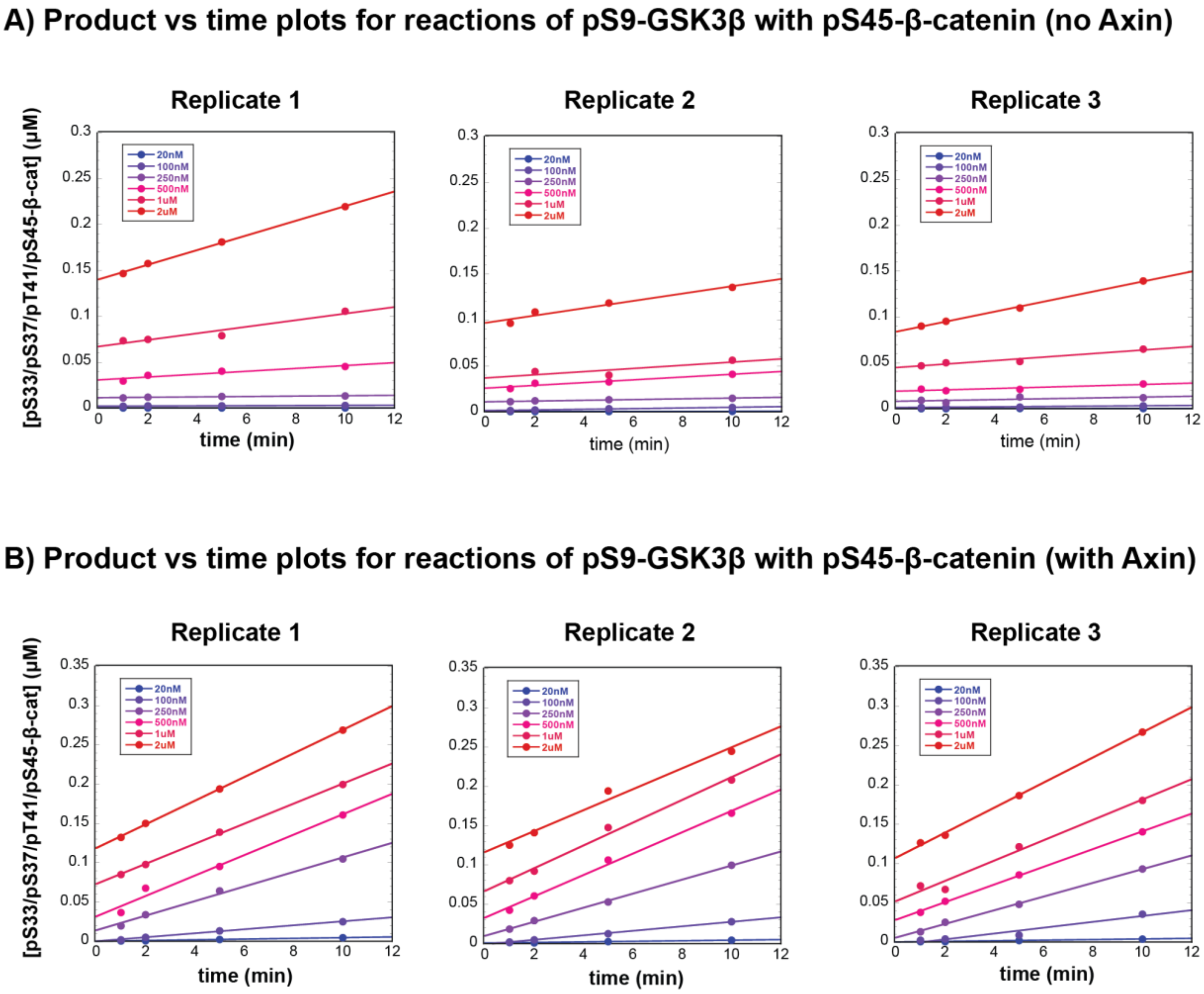


**Figure S7. Plots of product vs. time for reaction of pS9-GSK3β with pS45-β-catenin in the presence and absence of Axin, related to Figures 2 and 3.**
A) Product vs. time plots for reactions of pS9-GSK3β with pS45-β-catenin in the absence of Axin. B) Product vs. time plots for reactions of pS9-GSK3β with pS45-β-catenin in the presence of 500 nM Axin. Data in (A) and (B) correspond to the reaction conditions and western blots shown in Figure S6.

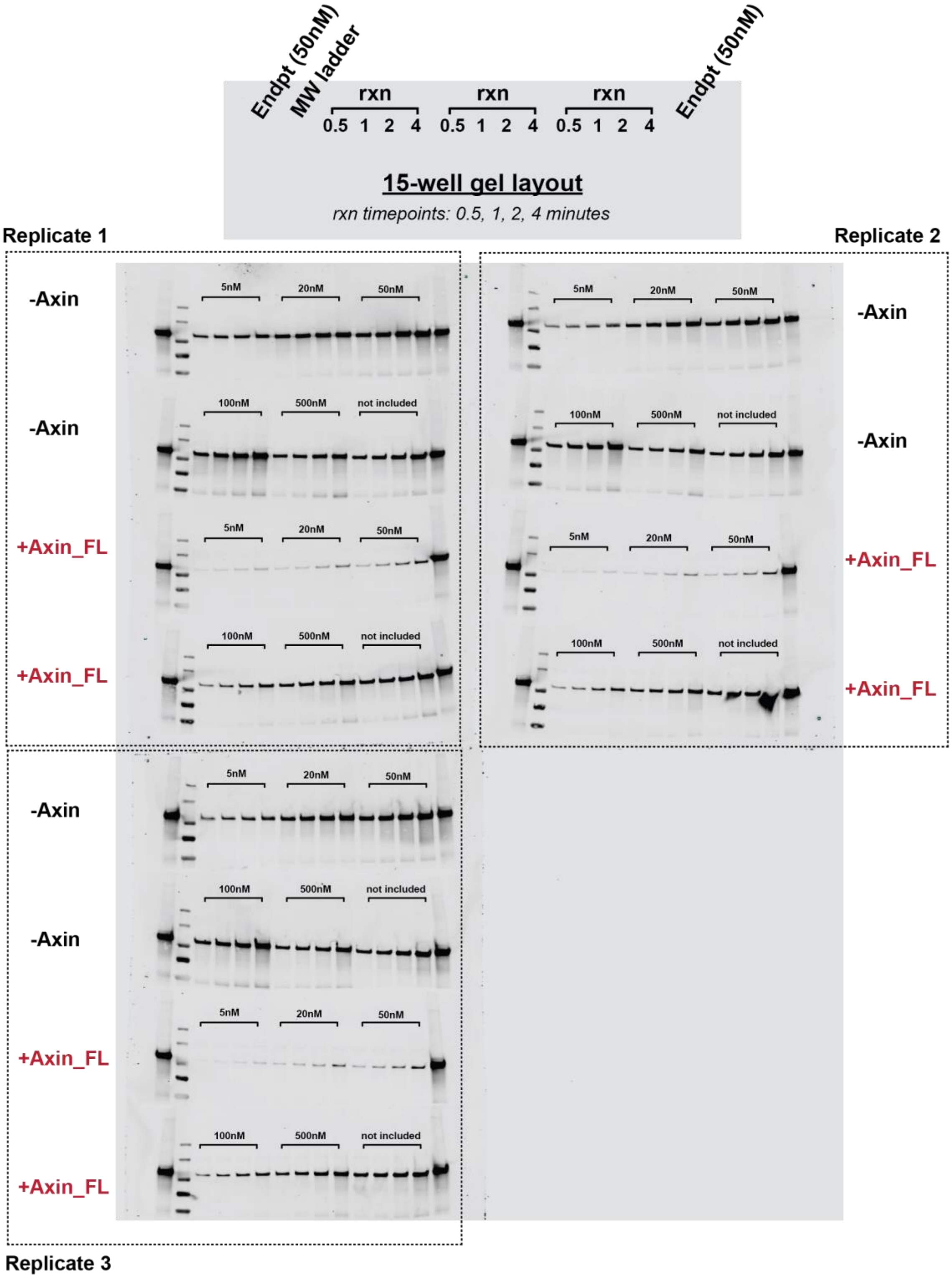


**Figure S8. Representative western blots for the reaction of PKA with GSK3β in the presence and absence of Axin, related to Figure 2.**

Western blots for reactions of varying concentrations of GSK3β with 20 nM PKA +/- 500 nM Axin. 500 nM GSK3β reaction gel samples were diluted 1:4 to prevent overloading the gel; all other reactions were diluted 1:2 (see Methods). See Figure S9 for quantification.

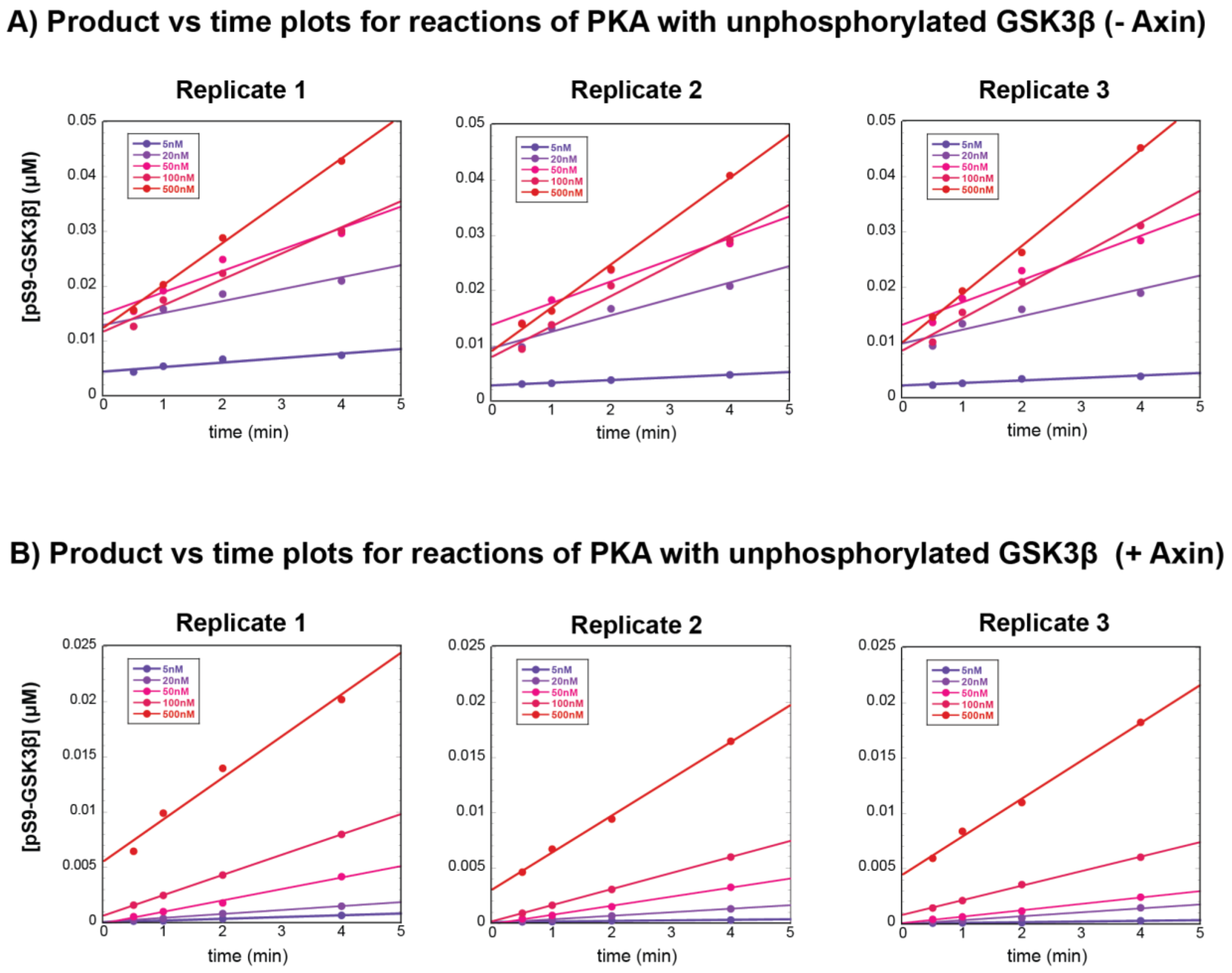


**Figure S9. Plots of product vs. time for reaction of PKA with GSK3β in the presence and absence of Axin, related to Figure 4.**

A) Product vs. time plots for reactions of PKA with GSK3β in the absence of Axin.

B) Product vs. time plots for reactions of PKA with GSK3β in the presence of 500 nM Axin. Data in (A) and (B) correspond to the reaction conditions and western blots shown in Figure S8.

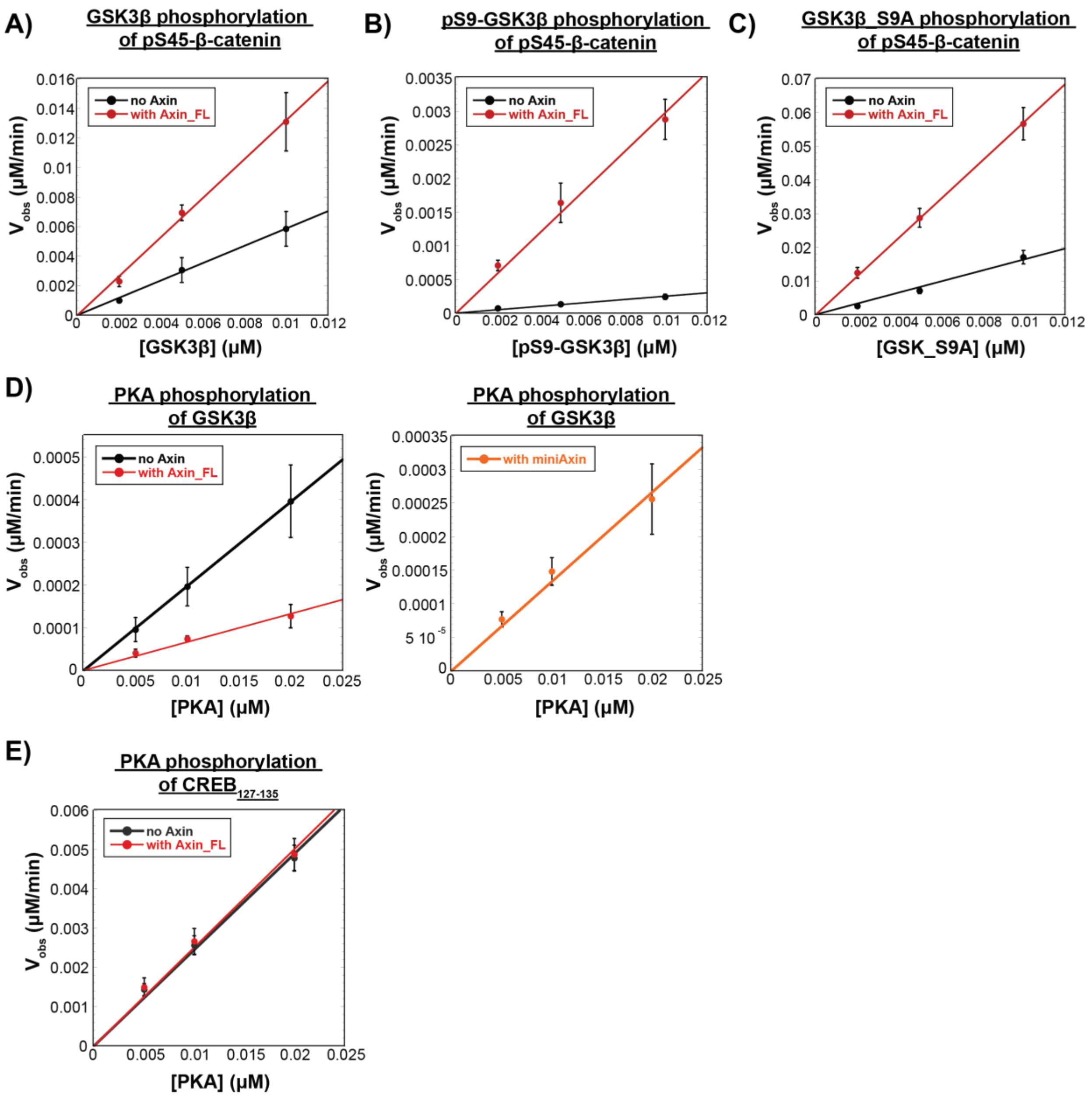


**Figure S10. $V_{obs}$ vs. [enzyme], related to Figure 2-4.**

$V_{obs}$ for fixed concentrations of substrate with varying concentrations of GSK3β or PKA in the presence or absence of Axin.

A) Plot of $V_{obs}$ vs. [unphosphorylated GSK3β], [pS9-GSK3β], or [GSK3β_S9A] with 50 nM pS45-β-catenin in the presence and absence of 500 nM Axin. $V_{obs}$ increases linearly with enzyme concentration, as expected.

B) Plot of $V_{obs}$ vs. [pS9-GSK3β] with 50 nM pS45-β-catenin in the presence and absence of 500 nM Axin.

C) Plot of $V_{obs}$ vs. [GSK3β_S9A] with 50 nM pS45-β-catenin in the presence and absence of 500 nM Axin. D) Plot of $V_{obs}$ vs. [PKA] with 20 nM GSK3β in the presence and absence of 500 nM Axin_FL or miniAxin. (E) Plot of $V_{obs}$ vs. [PKA] at 20 nM $CREB_{127\text{-}135}$ in the presence and absence of 500 nM Axin. Error bars are mean ± SD for at least 3 measurements.

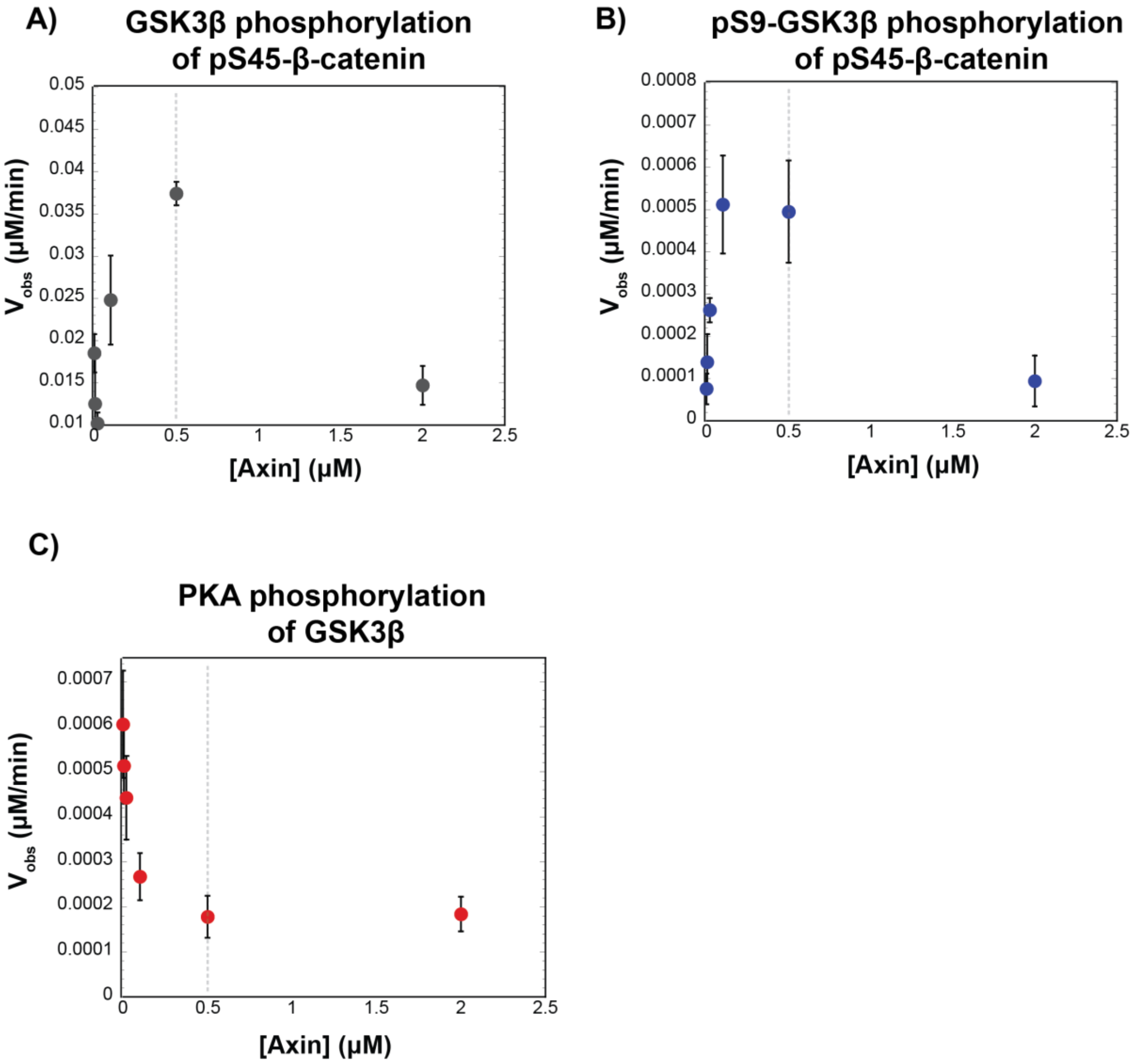


**Figure S11. Varying the concentration of Axin does not produce larger rate effects than observed with 500 nM Axin, related to Figures 2-4.**
A) Plots of $V_{obs}$ vs. [Axin] with 10 nM unphosphorylated GSK3β and 50 nM pS45-β-catenin
B) Plots of $V_{obs}$ vs. [Axin] with 10 nM pS9-GSK3β and 50 nM pS45-β-catenin.
C) Plots of $V_{obs}$ vs. [Axin] with 20 nM PKA and 20 nM GSK3β. Error bars are mean ± SD for at least 3 measurements.

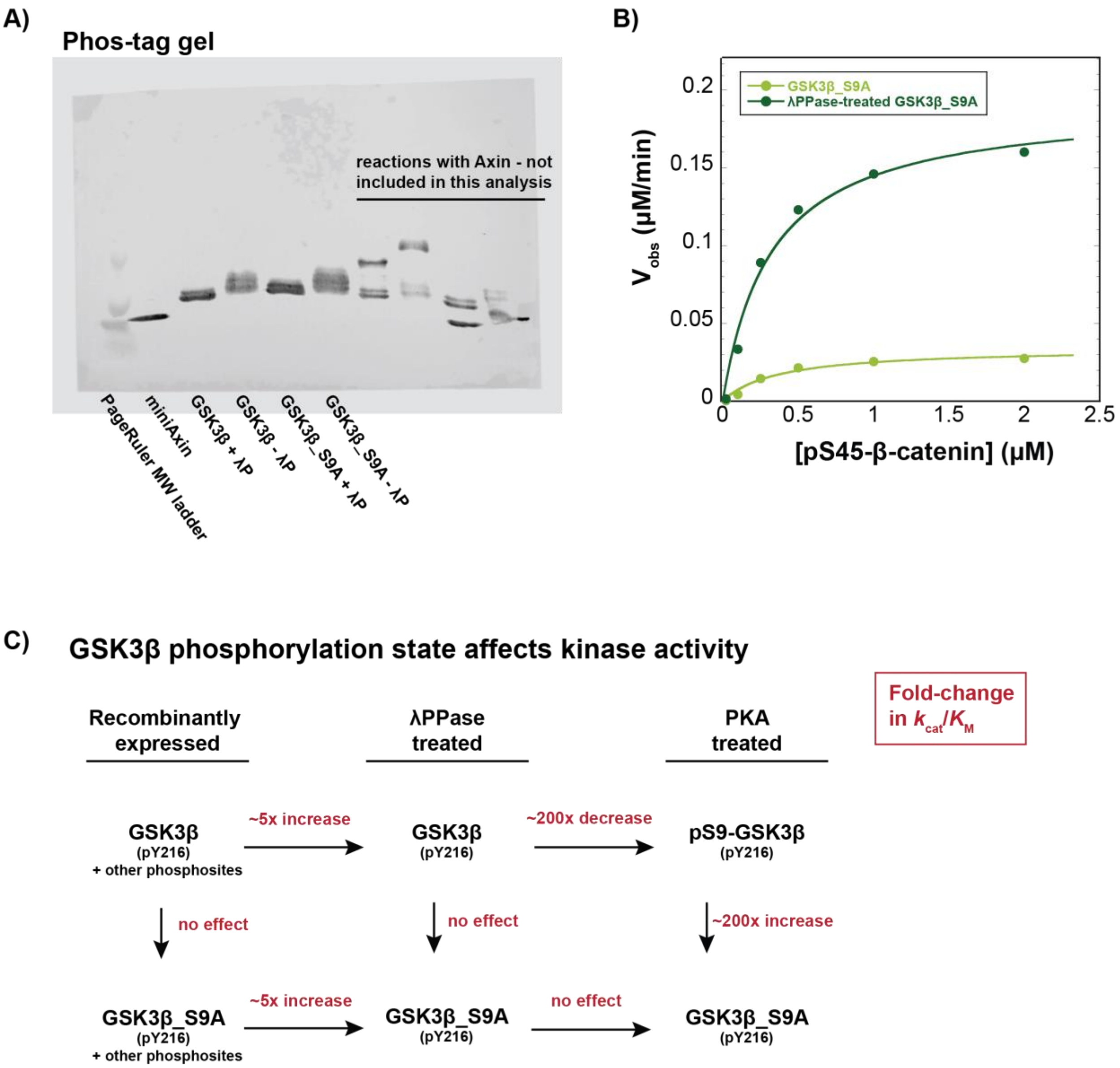
A)
Phos-tag gel
reactions with Axin - not included in this analysis
PageRuler MW ladder
miniAxin
GSK3β + λP
GSK3β - λP
GSK3β_S9A + λP
GSK3β_S9A - λP
B)
GSK3β_S9A
λPPase-treated GSK3β_S9A
V obs (μM/min)
[pS45-β-catenin] (μM)
0
0.05
0.1
0.15
0.2
0.5
1
1.5
2
2.5
C)
GSK3β phosphorylation state affects kinase activity
Recombinantly expressed
λPPase treated
PKA treated
Fold-change in kcat/KM
GSK3β (pY216) + other phosphosites
~5x increase
GSK3β (pY216)
~200x decrease
pS9-GSK3β (pY216)
no effect
no effect
~200x increase
GSK3β_S9A (pY216) + other phosphosites
~5x increase
GSK3β_S9A (pY216)
no effect
GSK3β_S9A (pY216)

**Figure S12. Recombinant GSK3β is phosphorylated on multiple sites.**
A) Recombinant GSK3β is phosphorylated at phosphosites other than Ser9. Phos-tag gel of GSK3β and GSK3β_S9A with and without treatment with lambda phosphatase. Phos-tag gels were prepared and run as previously described (Gavagan et al., 2020) (see Methods). Samples with GSK3β or GSK3β_S9A were prepared in PMP buffer (NEB) with 1 mM $MnCl_2$ and 400 nM GSK3β or GSK3β_S9A and incubated in the presence or absence of 30 μM lambda phosphatase for 30 min at 30 °C. The slower-migrating species are phosphorylated GSK3β or GSK3β_S9A. The presence of phosphorylated bands in GSK3β_S9A indicates that additional sites besides Ser9 are phosphorylated in recombinant GSK3β.
B) Michaelis-Menten plot of $V_{obs}$ versus [pS45-β-catenin] at 10 nM GSK3β_S9A or 10 nM lambda phosphatase-treated GSK3β_S9A. Dephosphorylation of GSK3β_S9A by lambda phosphatase produces an ~5-fold increase in $k_{cat}/K_M$, largely due to a ~5-fold increase in $k_{cat}$. See Table S4 for values of fitted kinetic parameters.
C) GSK3β kinase activity is affected by phosphorylation state at Ser9 and other phosphosites. Values on arrows are fold-change in $k_{cat}/K_M$ for different preparations of GSK3β in reactions with pS45-β-catenin. Dephosphorylation of wt GSK3β or GSK3β _S9A by lambda phosphatase produces ~5-fold increases in $k_{cat}/K_M$ (Table S5). PKA phosphorylation of lambda phosphatase-treated GSK3β to produce pS9-GSK3β leads to a ~200-fold decrease in $k_{cat}/K_M$ (Figure 2, Tables S1 and S5). PKA phosphorylation of lambda phosphatase-treated GSK3β_S9A produces no effect on $k_{cat}/K_M$ because GSK3β_S9A cannot be phosphorylated at the inhibitory Ser9 phosphosite (Figure 2, Table S5).

### Supplemental Tables

For all fitted kinetic parameters, see the Methods section for the kinetic model used to fit the data to obtain observed values of $k_{cat}$, $K_M$, and $k_{cat}/K_M$. Standard errors are from non-linear least squares fits to the initial rate data as described in the Methods.

**Table S1.** Kinetic parameters for GSK3β reactions with pS45-β-catenin, related to Figures 2-3.[a]

| Enzyme | Reaction | $k_{cat}$ ($s^{-1}$) | $K_M$ (μM) | $k_{cat}/K_M$ ($M^{-1}s^{-1}$) |
|---|---|---|---|---|
| pS9-GSK3β | –Axin | n.d. | n.d. (≥ 2 μM) | $(4.9 \pm 0.1) \times 10^3$ |
| | +full length Axin | $(2.9 \pm 0.3) \times 10^{-2}$ | 0.29 ± 0.08 | $(1.1 \pm 0.3) \times 10^5$ |
| GSK3β | –Axin | $(2.8 \pm 0.2) \times 10^{-1}$ | 0.33 ± 0.08 | $(8.5 \pm 2.1) \times 10^5$ |
| | +full length Axin | $(3.4 \pm 0.1) \times 10^{-1}$ | 0.17 ± 0.02 | $(2.0 \pm 0.2) \times 10^6$ |
| GSK3β_S9A | –Axin | $(2.8 \pm 0.2) \times 10^{-1}$ | 0.27 ± 0.05 | $(9.5 \pm 1.8) \times 10^5$ |
| | +full length Axin | $(3.3 \pm 0.1) \times 10^{-1}$ | 0.16 ± 0.02 | $(2.1 \pm 0.3) \times 10^6$ |

[a] See Figures 2 and 3 for data. The pS9-GSK3β reactions in the absence of Axin did not detectably saturate up to 2 μM substrate (Figures 2 and 3), and only the value of $k_{cat}/K_M$ could be accurately determined. No deviation from linearity was observed at 2 μM pS45-β-catenin (the highest pS45-β-catenin concentration tested), suggesting a conservative estimate that $K_M \geq 2$ μM. pS9-GSK3β, GSK3β, and GSK3β_S9A were coexpressed with lambda phosphatase. pS9-GSK3β and GSK3β_S9A were incubated with PKA and ATP before use (see Methods). The $k_{cat}/K_M$ for λPPase-treated GSK3β is ~5-fold higher than for non-λPPase-treated GSK3β used in previous studies (Gavagan et al., 2020) (see Figure S12C and Table S5).

**Table S2.** Kinetic parameters for PKA reactions, related to Figure 4.

| Substrate | Reaction | $k_{cat}$ ($s^{-1}$) | $K_M$ (μM) | $k_{cat}/K_M$ ($M^{-1}s^{-1}$) |
|---|---|---|---|---|
| GSK3β | –Axin | $(7.5 \pm 0.3) \times 10^{-3}$ | 0.062 ± 0.007 | $(1.2 \pm 0.1) \times 10^5$ |
| | +full length Axin | $(4.4 \pm 0.1) \times 10^{-3}$ | 0.25 ± 0.02 | $(1.7 \pm 0.1) \times 10^4$ |
| | +miniAxin | $(4.8 \pm 0.4) \times 10^{-3}$ | 0.23 ± 0.05 | $(2.1 \pm 0.5) \times 10^4$ |
| $CREB_{127\text{-}135}$ | –Axin | $(2.3 \pm 0.04) \times 10^{-2}$ | 0.15 ± 0.01 | $(1.5 \pm 0.1) \times 10^5$ |
| | +full length Axin | $(2.3 \pm 0.05) \times 10^{-2}$ | 0.14 ± 0.01 | $(1.6 \pm 0.1) \times 10^5$ |

**Table S3.** $K_{M,\ ATP}$ values for all reactions, related to Figure S3.

| Enzyme | Substrate | Reaction | $K_{M,\ ATP}$ (μM) |
|---|---|---|---|
| GSK3β | pS45-β-catenin | –Axin | 5.6 ± 0.9 |
| | | +full length Axin | 3.7 ± 1.3 |
| PKA | GSK3β | –Axin | 3.0 ± 0.6 |
| | | +full length Axin | 3.3 ± 0.8 |
| PKA | $CREB_{127\text{-}135}$ | –Axin | 2.1 ± 0.4 |
| | | +full length Axin | 2.1 ± 0.2 |

**Table S4.** Kinetic parameters for pS45-β-catenin reactions with non-PKA treated GSK3β_S9A with and without λPPase treatment, related to Figure S12.[a]

| Enzyme | Reaction | $k_{cat}$ ($s^{-1}$) | $K_M$ (μM) | $k_{cat}/K_M$ ($M^{-1}s^{-1}$) |
|---|---|---|---|---|
| GSK3β_S9A | Not treated | $(5.7 \pm 0.05) \times 10^{-2}$ | 0.37 ± 0.1 | $(1.5 \pm 0.5) \times 10^{5}$ |
| | λPPase-treated | $(3.2 \pm 0.2) \times 10^{-1}$ | 0.32 ± 0.1 | $(9.9 \pm 2.2) \times 10^{5}$ |

[a] λPPase-treated GSK3β_S9A was coexpressed with lambda phosphatase before use (see Methods). Untreated GSK3β_S9A was expressed without lambda phosphatase.

**Table S5.** Values of $k_{cat}/K_M$ for untreated, λPPase-treated, and PKA-treated GSK3β and GSK3β_S9A in reactions with the substrate pS45-β-catenin, related to Figure S12.[a]

| Enzyme | Reaction | $k_{cat}/K_M$ ($M^{-1}s^{-1}$) |
|---|---|---|
| GSK3β | Untreated | $(1.8 \pm 0.2) \times 10^{5}$ |
| | λPPase-treated | $(8.5 \pm 2.1) \times 10^{5}$ |
| | PKA-treated | $(4.9 \pm 0.1) \times 10^{3}$ |
| GSK3β_S9A | Untreated | $(1.5 \pm 0.5) \times 10^{5}$ |
| | λPPase-treated | $(9.9 \pm 2.2) \times 10^{5}$ |
| | PKA-treated | $(9.5 \pm 1.8) \times 10^{5}$ |

[a] See (Gavagan et al., 2020) and Figures 2, 3, 4, and S12 for data. λPPase-treated GSK3β and GSK3β_S9A were coexpressed with lambda phosphatase before use. PKA-treated GSK3β and GSK3β_S9A were coexpressed with lambda phosphatase and then incubated with PKA and ATP before use (see Methods). PKA-treated GSK3β is pS9-GSK3β. The $k_{cat}/K_M$ value for untreated GSK3β is from previous work (Gavagan et al., 2020).

**Table S6.** Protein expression plasmids, related to Methods

| Plasmid | Protein[a] | Expressed Protein | Vector[b] | Source |
|---|---|---|---|---|
| pMG024[c] | GSK3β | MBP-GSK3β-HA-His | pMBP-MG | *This study* |
| | Lambda phosphatase (λPPase) | GST_λPPase | | |
| pMG071[c] | GSK3β_S9A | MBP-GSK3β_S9A-HA-His | pMBP-MG | *This study* |
| | Lambda phosphatase (λPPase) | GST_λPPase | | |
| pES001 | GSK3β | MBP-GSK3β-HA-His | pMBP-MG | *(Gavagan et al., 2020)* |
| pES002 | GSK3β_S9A | MBP-GSK3β_S9A-HA-His | pMBP-MG | *This study* |
| pEF073 | Axin | MBP-Axin-His | pMBP-MG | *(Gavagan et al., 2020)* |
| pMG023 | $Axin_{384-518}$ (miniAxin) | MBP-$Axin_{384-518}$-His | pMBP-MG | *(Gavagan et al., 2020)* |
| pMG051[d] | β-catenin | MBP-β-catenin-His | pMBP-MG | *(Gavagan et al., 2020)* |
| | CK1α | GST-CK1α | | |
| pEF086 | CREB (127-135) | MBP-$CREB_{127-135}$-His | pMBP-MG | *(Gavagan et al., 2020)* |
| pMG026 | Lambda phosphatase (λPPase) | His-λPPase | pBH4 | *This study* |
| $H_6$-rC | PKA catalytic subunit | His-PKA-rC | pET15b | Addgene #14921 |

[a] All proteins are human sequences except PKA, which is the mouse sequence.

[b] pMBP-MG is a modified version of pMAL-p2X (New England Biolabs) with an N-terminal TEV-cleavable MBP tag and a C-terminal $His_6$ tag. pBH4 is a modified version of pET15b (Novagen) with an N-terminal TEV-cleavable $His_6$ tag. pBH4 and pMBP-MG were described previously (Good et al., 2009).

[c] pMG024 was constructed by inserting GST from pETARA (Good et al., 2009) and the λPPase expression cassette (without the His tag) from pMG026 into the pES001 backbone. The GSK3β_S9A mutant (pMG071) was cloned into this dual expression cassette.

[d] pMG051 was constructed by inserting the GST-CK1α expression cassette (without the His tag) from pMG046 into the pEF019 backbone.
